# Nutritional condition drives spatial variation in physiology of Antarctic lipid-storing copepods

**DOI:** 10.1101/2023.09.25.559317

**Authors:** Cory A. Berger, Deborah K. Steinberg, Ann M. Tarrant

## Abstract

Lipid-rich copepods form an essential link between primary producers and higher trophic levels in high-latitude oceans. These zooplankton can take advantage of ephemeral phytoplankton blooms to fuel development and reproduction. However, we have limited understanding of how the physiological condition of these animals varies in relation to environmental factors such as food availability. Due to high advection, it is likely that physiological plasticity, rather than local adaptation, is primarily responsible for physiological differences within a region. We use transcriptomics and other physiological metrics to understand how two species of copepods (*Calanoides acutus* and *Calanus propinquus*) vary across environmental gradients along the West Antarctic Peninsula. For the primarily herbivorous *C. acutus*, physiological separation between sampling locations appears to be driven by feeding status, and gene expression differences indicate differential expression of genes regulating lipid metabolism, reproduction, aerobic metabolism, and protein translation. For the more omnivorous *C. propinquus,* physiology and gene expression did not segregate as clearly by location, showed minimal signs of food deprivation at any location, and had a weaker relationship with chlorophyll compared to *C. acutus*. By comparing these results with concurrent starvation experiments, we find thatspatial variation in gene expression reflects short-term differences in food availability (particularly for *C. acutus,*), and we identify genes whose expression indicates recent feeding status. Further examination of the relationships between food availability, copepod physiology, and population dynamics will ultimately improve our capacity to predict how copepod populations will respond to rapidly changing environmental conditions in the West Antarctic Peninsula ecosystem.

## Introduction

In temperate and polar marine ecosystems, copepods in the family Calanidae are key consumers of phytoplankton and microzooplankton (Campbell et al., 2009; Froneman et al., 2000; Thibault et al., 1999). These copepods have evolved traits that enable them to optimize feeding, growth and reproduction during times of high food availability, and to enter dormancy (diapause) during times of low food availability (Baumgartner & Tarrant, 2017; Hirche, 1983). In polar ecosystems, diapause is used as an overwintering strategy, during which copepods migrate into deep water, slow their metabolism, and subsist on stored lipids (Kattner et al., 2007; Lee et al., 2006). Stored lipids are also a primary source of energy to support egg production for some species, and for species that depend on continued feeding to support reproduction, stored lipids can help buffer energy shortages during gaps in food availability (Sainmont et al., 2014). Due to this strategy of seasonal lipid accumulation, dormancy, and reproduction, these species can utilize phytoplankton blooms as a primary food source and are themselves a rich food source for predators, such as fish, seabirds and baleen whales.

Antarctic and sub-Antarctic calanid copepods exhibit particularly diverse strategies to optimize their use of highly variable food supplies in seasonal environments. *Calanoides acutus* relies heavily on large phytoplankton as a dietary source, and most individuals overwinter in diapause (Schnack-Schiel et al., 1991). In contrast, *Calanus propinquus* has a comparatively omnivorous diet, can overwinter in surface waters, and is only weakly dependent on diapause (Bathmann et al., 1993; Pasternak et al., 2001; Schnack-Schiel et al., 1991). This opportunistic strategy enables, *C. propinquus* to be abundant under sea ice during late winter and can obtain about half of its dietary carbon from ice algae-derived prey over short time scales (Kohlbach et al., 2016; Kohlbach et al., 2018). *C. acutus* and *C. propinquus* also differ in their primary form of lipid storage (Hagen et al., 1993; Kattner et al., 1994; Ward et al., 1996). *C. acutus* stores lipids primarily in the form of wax esters, similar to the Arctic *Calanus* species. In contrast, *C. propinquus* (and the sub-Antarctic *Calanus simillimus*) primarily stores triglycerides. Despite these differences, *C. acutus* and *C. propinquus* are both broadly distributed throughout the Southern Ocean and have similar spatio-temporal patterns of abundance at large scales (Marin, 1988). Within the surface waters of the West Antarctic Peninsula (WAP), the focal region for this study, copepods comprise the dominant component of the surface (0-300 m) mesozooplankton during summer. *C. acutus* is among the most abundant large copepods in this region, and *C. propinquus* is much less abundant, but comprises a substantial portion of the mesozooplankton biomass due to its large size (Gleiber, 2014).

The Southern Ocean and Antarctic coastal seas have been experiencing pervasive changes in physical conditions, including temperature, circulation, and ice cover. The WAP is among the most rapidly warming regions on Earth, experiencing a 2.8°C increase in average annual air temperature from the 1950s to 2000 (Turner et al., 2016) and a decrease in the extent and seasonal duration of sea ice comparable to the greatest rates observed in the Arctic (Stammerjohn & Maksym, 2017; Stammerjohn et al., 2012). Along with this, changes in the distribution and abundance of Antarctic organisms have been documented at all trophic levels (reviewed by Constable et al., 2014; Rogers et al., 2020; Turner et al., 2014). Phytoplankton productivity and community composition have shown large interannual and regional variability (Schofield et al., 2017; Venables et al., 2013), The southern region has shown a trend toward increasing frequency of conditions favoring a shallow seasonal mixed layer depth that is associated with large diatom blooms and increased primary production, whereas no trends are discernable in the northern region (Schofield et al., 2018; Schofield et al., 2017). The abundance of copepods has increased in association with sea ice retreat and elevated chlorophyll a (chl a; Gleiber, 2014). However, it is unknown how copepods will respond to predicted reductions in the abundance of large phytoplankton, an important food source for lipid-storing copepods. Predicting responses of copepods to future environmental changes, including changes in phytoplankton abundance, can be informed by examining physiological variation in relation to current variations in environmental conditions.

Gene expression profiling is increasingly being used to infer physiological conditions within natural populations. Among pre-adults of the copepod *Neocalanus flemigeri* in the Gulf of Alaska, correlations of metabolic gene expression with food availability suggested a substantial capacity for physiological plasticity in response to patchy food supply (Roncalli et al., 2019). In contrast, *Calanus glacialis* pre-adults in the eastern Bering Sea showed relatively uniform expression of metabolic biomarkers, suggesting that they had experienced favorable growth conditions throughout the study area (Tarrant, Eisner & Kimmel, 2021). While comparable molecular studies have not yet been conducted for Antarctic lipid-storing copepods, observations of egg production, lipid content and composition, and metabolic enzyme activity provide insight into their physiological condition. For example, within the marginal ice zone of the Weddell Sea, lipid content and citrate synthase activity in female *C. acutus* and *C. propinquus* were low at ice-covered stations and higher at ice-edge and ice-free stations experiencing phytoplankton bloom or post-bloom conditions (Geiger et al., 2001).

We assessed physiological variation in adult females from populations of *C. acutus* and *C. propinquus* across environmental gradients within the WAP using transcriptomics, biochemical measurements, and visual observations. We predicted that aspects of their metabolic and reproductive condition, particularly of the more herbivorous *C. acutus*, would correlate with environmental chlorophyll concentration, and that aspects of condition would also correlate with local abundance of these species. In addition, we compared the observed physiological variation with responses to experimental feeding manipulations conducted at the same time (Berger et al., 2023) to more directly link gene expression patterns observed in the field with responses to food availability, and to identify putative molecular markers of metabolic status in these species.

## Methods

### Sampling

Zooplankton were sampled during austral summer 2019 aboard the “ARSV *Laurence M. Gould*” in association with sampling conducted by the Palmer Antarctica Long-Term Ecological Research (PAL-LTER) program. Samples were collected within the PAL-LTER sampling grid, which is positioned along the WAP and across the continental shelf (Fig. 1A, Waters & Smith, 1992). Stations within the grid are designated by a pair of numbers indicating the distance along the shelf followed by the distance across the shelf (e.g., 600.000 is a northern, inshore station). At each station, hydrographic properties were profiled using a CTD (SeaBird Electronics Seacat SBE 19plus sensor), and chl *a* concentration was profiled using a WET Labs ECO-FL fluorometer.

**Figure 1:**
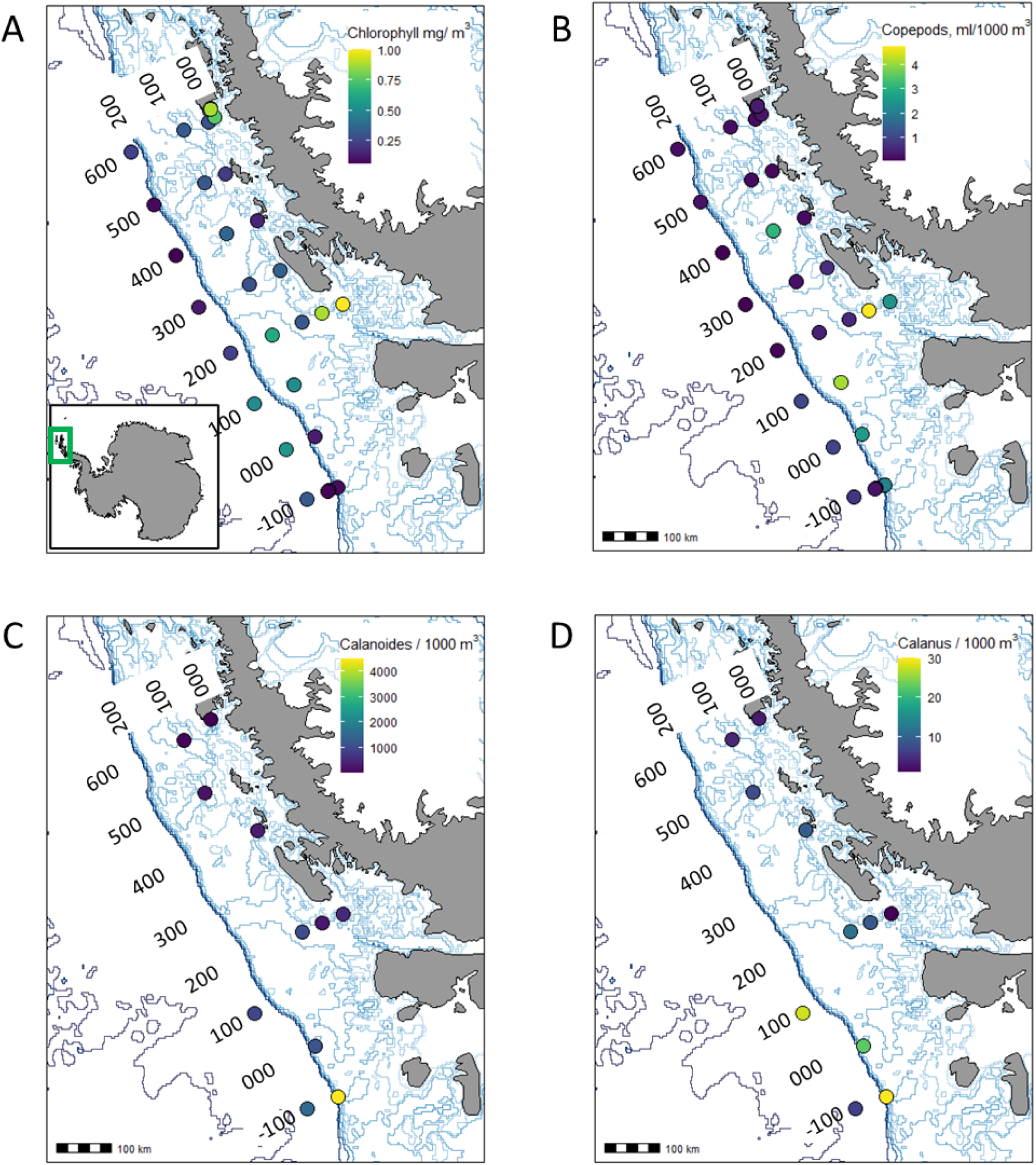
Spatial patterns in chlorophyll *a* concentration and copepod abundance along the Western Antarctic Peninsula in January 2019. Numbers indicate grid of station names (see Methods). Light blue lines indicate bathymetric contours, and heavy blue line indicates continental shelf break. (A) Mean chlorophyll *a* in the upper 200 m of the water column; green box in inset shows study area. (B) Total copepod biovolume (ml) per 1000 m^3^ seawater (sampled with 700 µm mesh net). (C-D) Abundance of *Calanoides acutus* (C) and *Calanus propinquus* (D) from a subset of stations sampled (with 333 µm mesh net), note difference in abundance scales.

Zooplankton were sampled via double oblique net tows with rectangular frame nets: macrozooplankton were sampled from 0-120 m using a 2 x 2 m net with 700-µm mesh, and mesozooplankton were sampled from 0-300 m using a 1 x 1 m net with 333-µm mesh. Aboard the ship, macrozooplankton samples were sorted taxonomically, and the biovolume and numbers of copepods and other taxa were quantified and whole or quantitative subsamples preserved in buffered formalin (Ross et al., 2008; Steinberg et al., 2015).

To sample copepods for physiological and transcriptomic studies, tows were conducted to a depth of either 120 or 200 m using the 700-µm mesh net. Copepods were contained in an ice-chilled bucket during sorting, adult female copepods were identified and individually photographed under a stereomicroscope. For transcriptional profiling, copepods were preserved in RNAlater at -20°C. For citrate synthase activity, copepods were stored at -80°C. Samples were shipped to Woods Hole Oceanographic Institution on dry ice. Because the distributions of *C. propinquus* and *C. acutus* differed and *C. propinquus* was less abundant, the numbers of stations analyzed and samples per station differed between species for each measurement type, as subsequently described.

### Copepod abundance and environmental correlations

*C. acutus* and *C. propinquus* were enumerated within mesozooplankton samples from 11 stations, including all the stations at which gene expression was characterized. Samples were split using a Folsom Plankton splitter. For each species a fraction ranging from 1/16 to the entire sample was counted, to either count at least 100 individuals or enumerate the entire sample. For these stations, correlations were examined among *C. acutus* and *C. propinquus* abundance, copepod abundance in the macrozooplankton samples, average temperature and chl *a* concentration in the upper 200 m (averaged from discrete measurements at each meter of depth), latitude, and bottom depth. A matrix of Spearman’s rank correlation coefficients was created using the rcorr function in the ‘hmisc’ R package. The strength and significance of correlations was visualized using the ‘corrplot’ R package (Wei & Simko, 2021).

### Citrate synthase activity

Copepods were thawed on ice, blotted on a lint-free tissue, and weighed on a Cahn C-33 microbalance. Groups of 2-5 copepods were pooled into 300 µL of ice-cold buffer (25 mM Tris, pH 7.8, 1mM EDTA, 10% glycerol) in a 5-mL Potter-Elvehjem homogenizer. Tissue was homogenized using a motorized PTFE pestle for two 30-second bursts with 30 seconds of ice cooling between bursts. Homogenates were centrifuged at 14,000g for 20 minutes at 4°C, and the supernatant was retained. Citrate synthase activity was measured with a modification the protocol of Hawkins et al. (2016) and normalized to wet weight, as described previously (Tarrant, McNamara-Bordewick, et al., 2021). Normality of the residuals and equality of variance were verified using the Shapiro Wilkes test and Levene’s tests, respectively. Differences among stations were tested using a t-test or one-way ANOVA; posthoc pairwise comparisons were tested using the holm method, as implemented in the ‘multcomp’ package.

### Photo analysis

Photographs were scored by two independent observers for the presence/absence of food in the gut and the degree of egg development. Egg development was scored in three categories ranging from 1 (no discernable eggs) to 3 (well-developed), using only photos in dorsal orientation for *C. acutus* and lateral orientation for *C. propinquus.* Photographs included copepods used for RNA-seq analysis and other copepods from the same tow. All photographs of sufficient quality were included in the analysis.

### RNA extraction, library construction and sequencing

Total RNA was extracted from individual copepods using the Aurum Fatty and Fibrous Tissue Kit (Bio-Rad) without DNase treatment. RNA yield and purity were measured using a Nanodrop Spectrophotometer. RNA was submitted to Arraystar Inc (Rockville, MD) for library construction and sequencing. Libraries were constructed using the KAPA Stranded RNA-seq Library Preparation Kit, and bar-coded libraries were sequenced with 150 base pair paired-end reads to a depth of 40 M reads on an Illumina HiSeq 4000. For *C. acutus,* libraries were constructed from individual samples; 5 libraries were sequenced from each of 6 stations. For *C. propinquus,* equivalent amounts of RNA from two individuals from the same station were pooled to create a sample for library construction, RNA-seq was conducted on 4 libraries from each of 4 stations. Sequence quality was assessed using FASTQC (v0.11.7, Babraham Bioinformatics).

#### Expression

Trimmed reads were mapped to the reference transcriptome for the appropriate species (Berger et al., 2023) using Salmon v1.1.0 (Patro et al., 2017) with the ‘gcBias’, ‘seqBias’ and ‘validateMappings’ flags and summarized to the gene level using the R package ‘tximport’ v1.20.0 (Soneson et al., 2015) with the ‘countsFromAbundance = lengthScaledTPM’ option. The gene expression matrix of the field samples (i.e., excluding the experimental samples from Berger et al., 2023) was filtered to retain clusters with at least 15 counts in at least 4 libraries. This retained 49,025/110,415 clusters for *C. acutus* and 53,379/143,667 for *C. propinquus* (Supplemental Files 2 and 3).

Sample distances were visualized using principal components analysis with the ‘prcomp’ R function. The PCA was performed on log-scaled counts per million (CPM) of all genes expressed above the threshold cutoff. Differential expression (DE) analysis was performed using ‘limma’ v3.48.3 (Phipson et al., 2016) with quality weights (Liu et al., 2015). For *C. acutus*, contrasts were performed between the two “station groups” as described in the Results; for *C. propinquus*, contrasts were performed between each station and the mean of the other three stations. Genes were considered DE with an adjusted p-value below 0.05. Weighted gene co-expression network analysis (WGCNA) was conducted using the combined set of field-collected and experimental samples as described in Berger et al. (2023). For *C. acutus*, a linear model (‘lm’ function in R) was used to compare module expression between station groups, and Pearson correlations were calculated between eigengene expression and environmental variables (log10 chl, mean temperature, and log10 abundance of *C. acutus*). For *C. propinquus*, module expression was compared across stations as in the DE analysis, but not with environmental variables because we judged that 4 field stations was too few to observe meaningful associations.

#### GO enrichment

Gene ontology (GO) terms that were enriched among DE genes (DEGs) or WGCNA modules were identified using GO_MWU (Wright et al., 2015) with genes ranked by log2(fold change). Semantically similar terms were summarized using Revigo (Supek et al., 2011) with the “Small” setting (0.5 SimRel similarity threshold). To explore differences among stations in energetic metabolism, reproductive physiology, and stress responses, we explicitly examined terms associated with *lipid metabolism* (GO:0006629), *protein metabolism* (GO:0019538), *reproduction* (GO:0000003) and *response to stress* (GO:0006950); descendant terms were retrieved using the R package GO.db v3.14.0.

#### Comparison with experimental feeding manipulations

Gene expression patterns in field-collected animals were compared with animals that were incubated aboard the ship for up to 9 days in fed or starved conditions (Berger et al., 2023). Log2(fold change) associated with field chl *a* measurements was correlated with log2(fold change) in the experimental response to starvation. We used discriminant analysis of principal components (DAPC) in the ‘adegenet’ package v2.1.8 (Jombart, 2008) to test whether the discriminant axis between fed sand starved animals also separated field samples. To identify possible biomarkers of feeding status, we selected genes with the most significant changes of expression associated apparently favorable feeding conditions in the field (stations 200.000 and 200.040; see Results) and experimental starvation. Putative biomarkers were required to have a minimum mean expression of 5 TPM10K, a log2(fold change) of at least 1 in the starvation experiment and the field comparison, and an unadjusted p-value of 0.001 in both tests. TPM10K is an expression measure similar to transcripts-per-million (TPM) except it also normalizes for the size of the transcriptome, and is thus more comparable between species (Munro et al., 2022).

## Results

### Environmental measurements and species distributions

Hydrographic profiling and zooplankton sampling were conducted at a grid of stations along the WAP in January 2019. Temperatures within the upper 200 m ranged from -1.84°C to 1.76°C, with the warmest maximum values in the outer slope stations and the warmest average temperatures in the northern inshore stations (Fig. S1). Chl *a* was highest at inshore stations along the 600 and 200 lines (Fig. 1A). Moderately high chl *a* was observed in the southern half of the grid along the shelf and in slope waters. Physiological and transcriptomic measurements were made using *C. acutus* and/or *C. propinquus* sampled from a total of 7 focal stations (6 stations *C. acutus*, 4 *C. propinquus*, 3 shared; Table S1). Among these stations, a northern inshore station (616.040) was characterized by warm water (>0.5 °C) throughout the upper 200 m and a strong surface chl *a* signal (Fig. 2). Three stations (200.000, 100.180 and 000.100) had shallow (<50 m) mixed layers and pronounced thermoclines, with the warmest surface water at the slope station (100.180). The other stations were more deeply mixed and had cold surface water. Substantial subsurface chl *a* maxima (peak concentration > 4 mg/m^3^) were present at 200.000 and 100.180, while 200.040 had moderately high chlorophyll (peak concentration 2.4 mg/m^3^), and the other stations had lower chl *a* concentrations throughout the water column.

**Figure 2:**
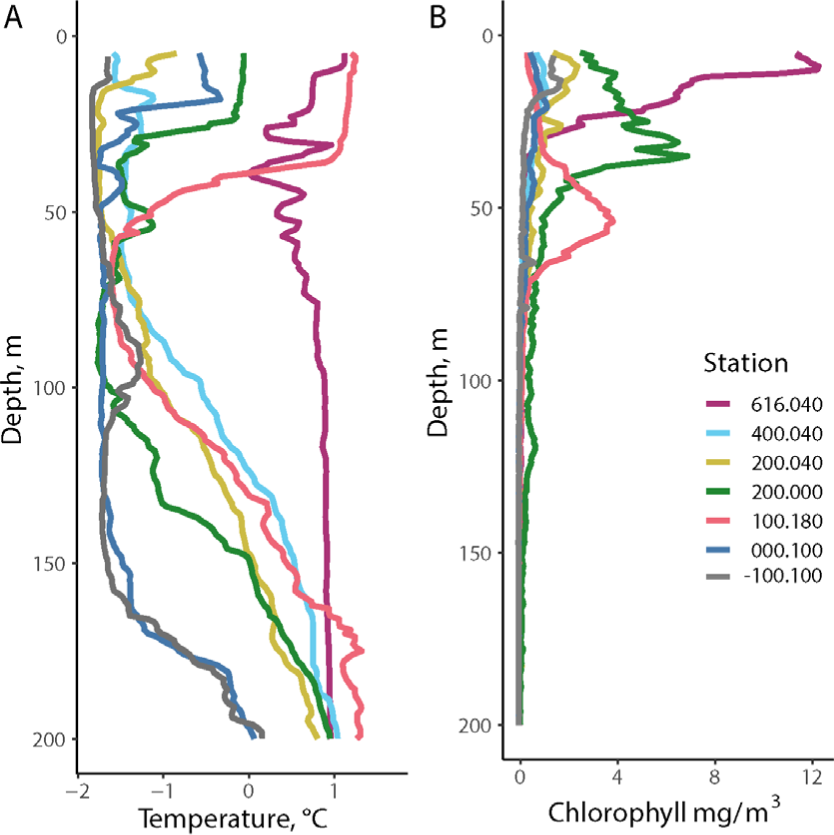
Depth profiles of (A) Temperature and (B) chlorophyll *a* for stations sampled for physiological measurements.

Overall abundance of large copepods was greatest at inshore, southern stations (Fig. 1B). *C. acutus* and *C. propinquus* abundance was determined at 11 stations, including all the focal stations (Figs. 1C-D; Table S1). As expected, *C. acutus* was much more abundant than *C. propinquus.* Both species were more abundant at the southern stations, and the highest abundance for both was observed at a far southern station (-100.100) that was near the shelf break and partially covered with ice. Abundances of *C. acutus* and *C. propinquus* were positively correlated with one another (r=0.62, p=0.04; Fig. 3). *C. acutus* was more abundant at cooler, high-latitude stations and *C. propinquus* was more abundant at cooler, low-chl *a* stations. Total copepod abundance within the mesozooplankton sample was positively correlated with latitude (r=0.71, p=0.02) but not with either of the focal species or any environmental factors.

**Figure 3:**
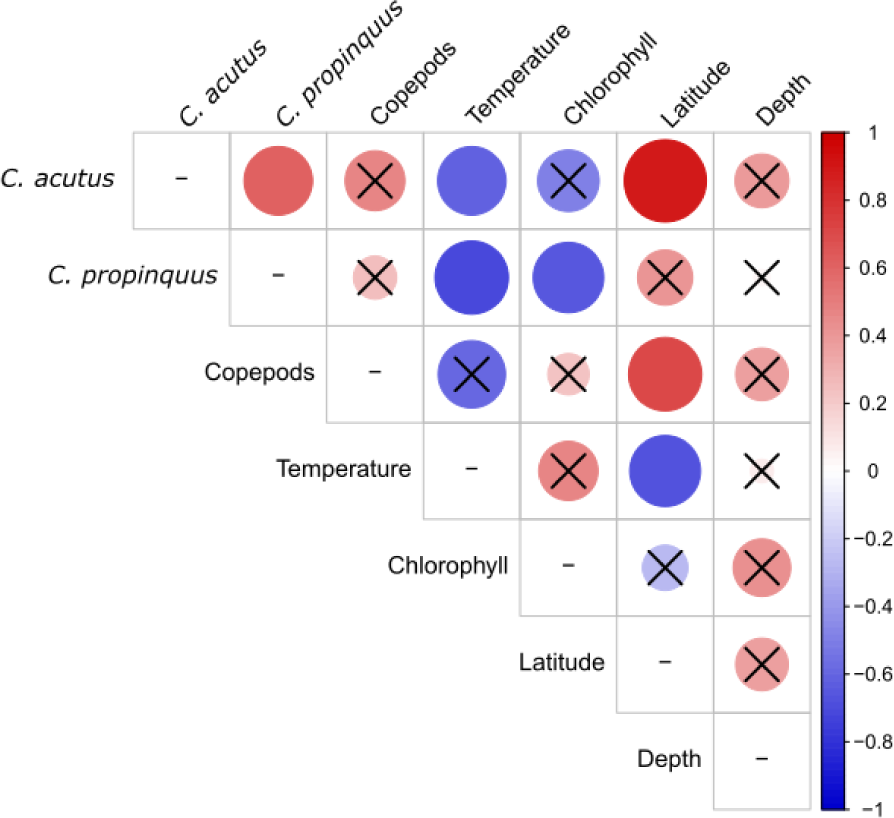
Heatmap showing Spearman’s correlations between pairs of measurements of copepod abundance and environmental parameters. From left to right: *Calanoides acutus* abundance*, Calanus propinquus* abundance, total copepod biovolume, average upper water column (0-200m) temperature, average (0-200m) chlorophyll *a* concentration, latitude, and water depth. Black crosses indicate statistically insignificant comparisons (p>0.05). Color scale and relative size both indicate correlation coefficient sign and magnitude.

### Physiological variation of *Calanoides acutus*

Principal component analysis (PCA) of *C. acutus* transcriptomic data revealed a pattern separating two mid-latitude inshore stations in Marguerite Bay (200.040 and 200.000, subsequently “200.xxx”) from all others (Fig. 4A), referred to as “station groups” hereafter. The 200.xxx stations exhibited intermediate temperatures and moderate-to-high chl *a* levels when averaged over the upper 200 m, from which the copepods were sampled. Station 200.000 had a warm (∼0°C) surface layer, a pronounced thermocline around 25 m, and a strong chl *a* peak below the thermocline. Station 200.040 was partially ice-covered, had colder and less stratified surficial water, and moderately high chl *a* that peaked near the surface. *C. acutus* abundance at these stations was low relative to the southern stations.

**Figure 4:**
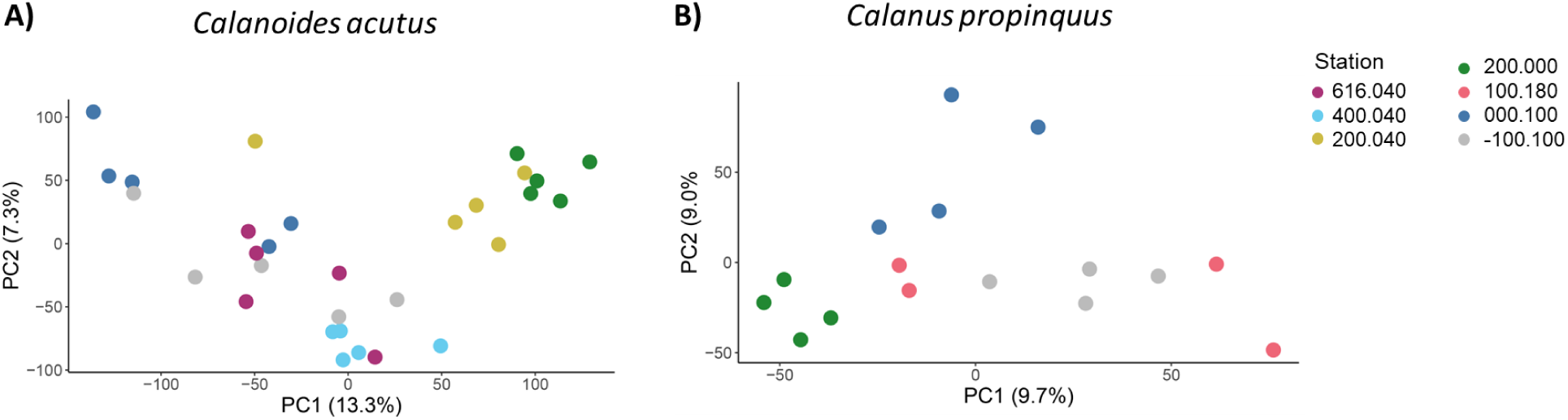
Overview of *Calanoides acutus* and *Calanus propinquus* gene expression patterns by station. Principal component analysis of (A) *C. acutus* and (B) *C. propinquus* gene expression. Axes show percent variation in gene expression explained by each principal component. Colors indicate field stations.

Food was visible within the guts of 35% of copepods from 200.xxx (15 of 43) but not in copepods from any of the other stations (0 of 35; Χ^2^, p < 0.01; Fig. 5A; Table S2). A higher proportion of copepods had well-developed eggs within 200.xxx (25%) vs. other stations (15%), but this difference was not significant (Fig. 5C; Table S2). Citrate synthase activity was higher in copepods from 200.xxx than in the two other stations for which measurements are available (ANOVA with Holm post-hoc test, p < 0.05; Fig. 5E; Table S3).

**Figure 5:**
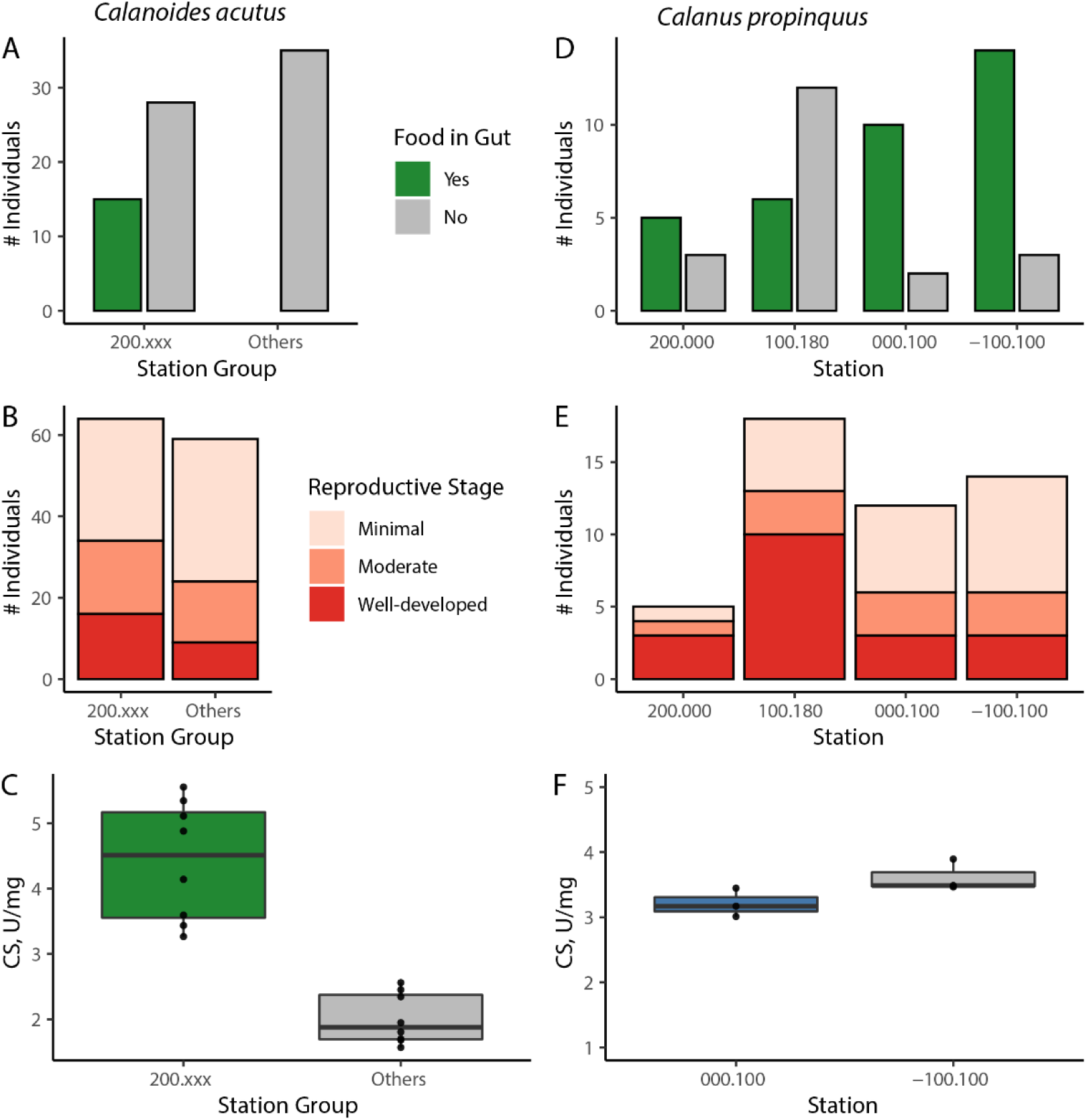
Physiological characteristics of *Calanoides acutus* (left) and *Calanus propinquus* (right). *C. acutus* data pooled by station group (see text). A, D: number of individuals having or lacking food in gut (in A no animals from “Other” stations had food in gut); B, E: relative egg development; C, F: Citrate synthase activity, Units per mg wet weight. The line in the box indicates the median, and the box encompasses the 25^th^ through 75^th^ percentiles of the data.

A pairwise contrast between station groups identified 12,043 differentially expressed genes (DEGs, adj. p. value < 0.05; Table 1), with 8712 genes upregulated at stations 200.xxx and 3341 downregulated.

**Table 1:** Differentially expressed genes between *Calanoides acutus* station groups (200.000 and 200.040 = 200.xxx vs all other stations)

|  | All | Annotated | Up Fed<sup>a</sup> | Up Starve<sup>a</sup> |
| --- | --- | --- | --- | --- |
| Up 200.xxx | 8712 | 4051 | 4 | 1674 |
| Down 200.xxx | 3341 | 2507 | 846 | 3 |

<sup>a</sup> Genes that were also differentially expressed in an associated feeding experiment (Berger et al., 2023)

Genes with higher expression at stations 200.xxx were enriched for gene ontology (GO) terms related to lipid metabolism, protein metabolism, reproduction and response to stress (Table S4, Supp File 2). Upregulated genes within *regulation of lipid metabolic process* included genes with conserved roles in regulating lipid homeostasis, such as an E75-like nuclear receptor, and NF2L1 homolog, (Cáceres et al., 2011; Hirotsu et al., 2012), but also a zinc finger protein, prosaposin and alkaline ceramidase, which have less certain roles in metabolic regulation. Among the stress-related genes, *response to hypoxia* was enriched. Among the 104 upregulated genes associated with this term were hypoxia inducible factor1A, catalase, soluble guanylate cyclases, and 85 genes annotated as “REI-silencing transcription factor”. Genes with lower expression at stations 200.xxx, were enriched for GO terms associated with protein metabolism (*cytoplasmic translation, regulation of translation, proteolysis, protein ubiquitination, and mitochondrial translation*) and stress response (*DNA repair* and *cellular response to stress*). No enriched terms were associated with lipid metabolism or reproduction.

The list of DEGs was inspected for genes previously associated with lipid synthesis and reproduction in *Calanus spp,* as well as the DEGs with the with largest fold-changes. Very-long chain fatty acid elongases (ELOV/ELVL), fatty acid desaturases (also called acyl-CoA desaturases, “desaturases” hereafter) and fatty acid binding proteins (FABPs) are involved in the synthesis and transport of storage lipids; these groups are subsequently referred to as “lipid-associated genes”. Many of these genes were upregulated at stations 200.xxx (12 ELOV/ELVLs, 13 desaturases, and 17 FABPs), and only a single FABP was downregulated at these stations.

Three desaturases were among the most strongly upregulated genes. Genes annotated as digestive lipases (gastric or pancreatic lipase) showed both directions of regulation, but one “gastric triacylglycerol lipase” was strongly upregulated. Vitellogenins are lipoproteins that are major components of egg yolk. Six vitellogenins were upregulated at stations 200.xxx; none were downregulated. Genes that were strongly upregulated at 200.xxx relative to other stations included five ubiquitin oxidoreductases, a cytochrome c oxidase, and ATP synthase. Genes that were strongly downregulated at 200.xxx included FTZ-F1, Fem1, arginine kinase, a DNAJ family member, 2 phenoloxidase activating factors, a cytochrome P450 family4c3 homolog, and thioredoxin.

Weighted gene co-expression network analysis identified 25 groups of genes (modules) with highly correlated expression across a combined dataset of the field samples described here and samples from an associated shipboard experiment (Berger et al., 2023). Module eigengenes were associated with environmental conditions through Pearson correlations, and with station groups with a linear model (Fig. 6). As described below, we identified modules for which expression was correlated with station group and/or chl *a* concentration. No modules were correlated with either temperature or *C. acutus* abundance.

**Figure 6.**
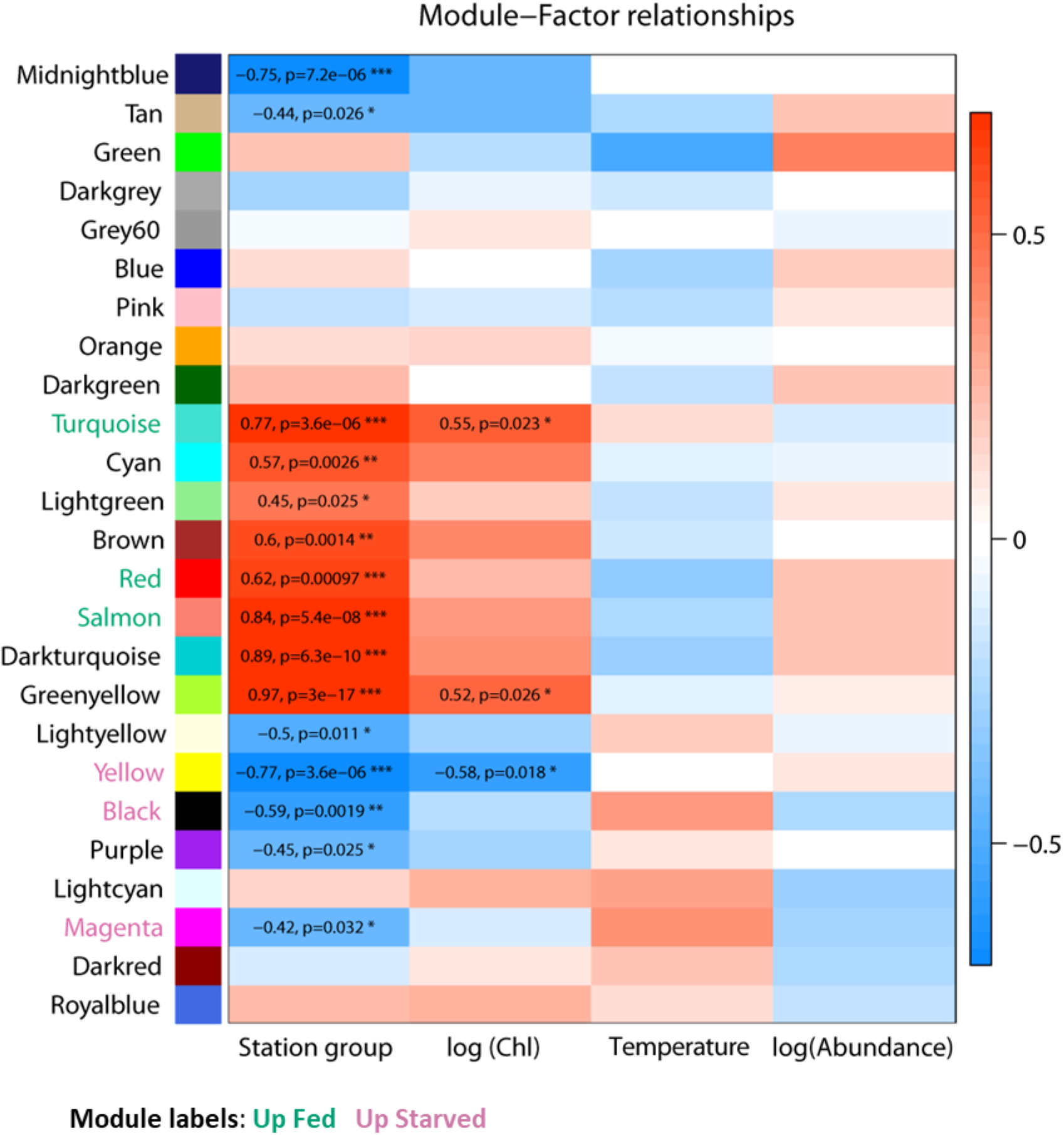
Associations of *Calanoides acutus* gene expression with collection stations, ecological factors and feeding history. Heatmap showing WGCNA module eigengene expression associated with collection station and ecological factors (log integrated Chl *a* in upper 200 m, average water temperature upper 200 m, log *C. acutus* abundance). Model labels are colored to indicate comparison with results from a shipboard feeding experiment (see also Berger et al., 2023). Eigengene expression was regressed against factor values. Colors represent the correlation coefficient and direction of regulation: red indicates positive correlations or upregulation at station group 200.xxx, blue negative correlations or downregulation at station group 200.xxx. Values of significant correlation coefficients are shown, along with the adjusted p-value from the linear model (*, p < 0.05; **, p < 0.01, ***, p < 0.001).

Eigengenes of 8 modules had higher expression at stations 200.xxx relative to the remaining stations (p<0.05). Of the 4051 upregulated and annotated genes, 85% were assigned to one of three modules: Turquoise, Greenyellow, or Red; the Turquoise and Greenyellow modules were also positively correlated with chl *a*. The Turquoise module was the largest module; in the station-group comparison, it included 2938 upregulated annotated genes, 11 lipid-associated genes, and all the differentially expressed vitellogenins. GO terms enriched in the Turquoise module corresponded to diverse processes including DNA metabolism, regulation and methylation; regulation of reproductive processes; cellular responses to stimuli; and muscle structure development (see Berger et al., 2023 for full GO enrichment results of WGCNA modules). The Greenyellow module included two lipid-associated genes and was enriched for GO terms related to translation and biosynthesis of amides/organonitrogen compounds. The Red module included 14 lipid-associated genes and was enriched GO for terms related to lipid and fatty acid synthesis, ion transmembrane transport, and amino acid transport and metabolism.

Eigengenes of 7 modules were downregulated in stations 200.xxx relative to the other stations. Of the 2507 downregulated and annotated genes, 74% were assigned to one of three modules: Yellow, Black or Magenta; the Yellow module was also negatively correlated with chl *a*. The Yellow module contained over half of the downregulated genes with annotation (1309 genes) and was associated with enriched GO terms related to RNA processing and metabolism, synthesis and catabolism of proteins and other macromolecules, mitochondrial respiratory chain assembly and aerobic respiration. The Black module was enriched with terms related to cell division, RNA processing and protein ubiquitination. The Magenta module was enriched for terms related to DNA replication and repair, cellular response to stress, intracellular transport, and RNA metabolism.

### Physiological variation of *Calanus propinquus*

For *C. propinquus,* transcriptomic data from the four stations sampled clustered into three groups: one with samples from 200.000, a second with samples from 000.100, and a third with samples from both 100.180 and -100.100 (Fig. 4B). Stations 100.180 and -100.100 had the highest *C. propinquus* abundances (Fig. 1D), but they did not share other similar environmental characteristics. Station 100.180 had warm surface water, a pronounced thermocline, and a strong subsurface chl *a* peak. In contrast, Station -100.100 was cold and well-mixed with relatively low chl *a* (Fig. 2).

Physiological observations did not provide any consistent basis for grouping stations. Food was visible within the guts of most individuals from 3 of the 4 stations (29 of 37; 63-83% per station), but a minority of individuals from 000.100 (6 of 18, 33%; Fig 5B; Table S5). Station 100.180 had the highest proportion of copepods with well-developed eggs (56% vs. 20-25% in other stations; Fig. 5D; Table S5). Due to limited specimen availability, citrate synthase activity was only measured at two stations; though the mean value was higher at -100.100 than at 000.100, this difference was not significant (t-test, t = -2.1547, df = 4, p-value = 0.09747; Fig. 5F).

Because gene expression and physiological observations did not provide a consistent basis for grouping stations, gene expression from each station was compared with the mean expression from the other three stations (Table 3; Supp. File 3). Relatively few DEGs were identified in any comparison, with the largest sets associated with 000.100 (282 upregulated, 94 downregulated) and 200.000 (137 upregulated, 35 downregulated). Among these, differential expression of lipid-associated genes and vitellogenin was only observed at 200.000, where we observed upregulation of 4 elongases, a desaturase and 2 vitellogenins (Table 2). This result is qualitatively similar to *C. acutus,* for which we observed upregulation of several lipid-associated genes and vitellogenins at Stations 200.xxx.

**Table 2:** Differentially expressed lipid-associated genes and vitellogenins in the copepods *Calanoides acutus* and *Calanus propinquus*.

| Species | *Calanoides acutus* |  | *Calanus propinquus* |  |
| --- | --- | --- | --- | --- |
| Contrast/Gene Category | Up 200.xxx | Down 200.xxx | Up 200.000 | Down 200.000 |
| Elongase (ELOV or ELVL) | 12 | 0 | 4 | 0 |
| Desaturase (Acyl-CoA or fatty acid) | 13 | 0 | 1 | 0 |
| Fatty acid-binding protein (FABP) | 15 | 3 | 0 | 0 |
| Vitellogenin or Vitellogenin receptor | 6 | 0 | 2 | 0 |

**Table 3:** Differentially expressed genes among *Calanus propinquus* stations.

| Station | Up<sup>a</sup> | Down<sup>a</sup> |
| --- | --- | --- |
| 200.000 | 137 | 3 |
| 100.180 | 22 | 0 |
| 000.100 | 282 | 94 |
| -100.100 | 36 | 4 |

<sup>a</sup> Expression at each station was compared with mean expression across all other stations.

As with *C. acutus*, we identified enriched GO terms related to lipid metabolism, protein metabolism, reproduction, and response to stress (Table S6). GO terms associated with higher gene expression at 200.000 included those related to dietary lipid metabolism, regulation of histone methylation, and meiosis. These processes were similarly regulated in *C. acutus* at Stations 200.xxx, though the specific enriched GO terms were not necessarily the same. These gene expression patterns may suggest favorable feeding conditions at 200.000. Other patterns in *C. propinquus* included an apparent upregulation of translation-related genes at 000.100 and downregulation at 200.000, as well as downregulation of reproductive processes and several stress-related processes at -100.000.

Linear models were used to associate WGCNA eigengenes with each site compared to the mean of all other sites (Fig. 7). Consistent with the relatively large number of DEGs, the largest number of modules (8) was associated with 000.100. Upregulated modules were enriched for GO terms related to nucleolus/RNP complexes (Lightgreen module), translation (Darkturquoise) and RNA processing and protein folding (Salmon). Downregulated modules were enriched for terms related to ribosomes and mitochondria (Greenyellow), chromatin replication (Lightyellow), triglyceride metabolism and ion transport (Midnightblue) and carboxlic acid, amino acid and carbohydrate metabolism (Royalblue).

**Figure 7.**
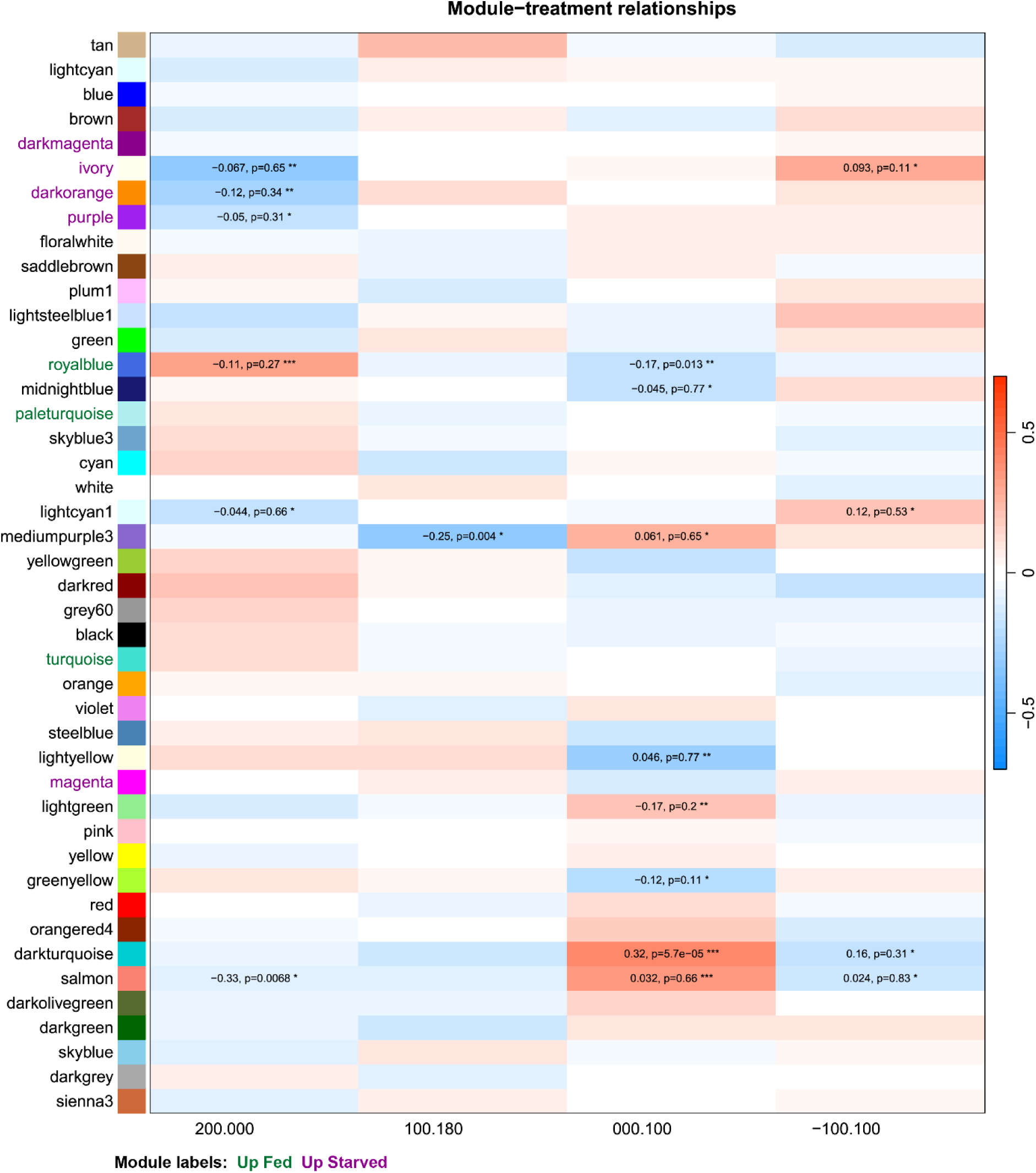
Associations of *Calanus propinquus* gene expression with collection stations. Heatmap showing WGCNA module eigengene expression associated with collection station. Model labels are colored to indicate comparison with results from a shipboard feeding experiment (see also Berger et al., 2023). Eigengene expression was regressed against factor values. The color represents the correlation coefficient and direction of regulation: red indicates upregulation relative to the mean expression across all other stations, and blue indicates downregulation. For significant correlations, the value is shown along with the adjusted p-value from the linear model (*, p < 0.05; **, p < 0.01, ***, p < 0.001).

### Correlation of field chlorophyll *a* measurements and experimental starvation response

In an accompanying paper, we measured gene expression changes in *C. acutus* and *C. propinquus* in response to short-term (9 days) starvation (Berger et al., 2023). In both species, the fold-change of genes that were DE in starved animals was correlated with the fold-change associated with field chl *a* in the present study (Spearman’s ρ = 0.73 in *C. acutus*, ρ = 0.44 in *C. propinquus*), suggesting that transcriptomic differences between high- and low-chl *a* stations might recapitulate the starvation response. To investigate this, we used discriminant analysis of principal components (DAPC) to examine whether field populations separated along the first discriminant axis between fed and starved experimental samples. This was the case for *C. acutus*, as low-chl *a* stations overlapped with or were close to the starved samples, and high-chl *a* sites overlapped with fed samples (Fig. 8A; the exception being Site 616.040; see Discussion). However, all field *C. propinquus* samples were close to the fed samples along the first discriminant axis, consistent with our observations that most *C. propinquus* had food in the gut (Fig. 8B).

**Figure 8.**
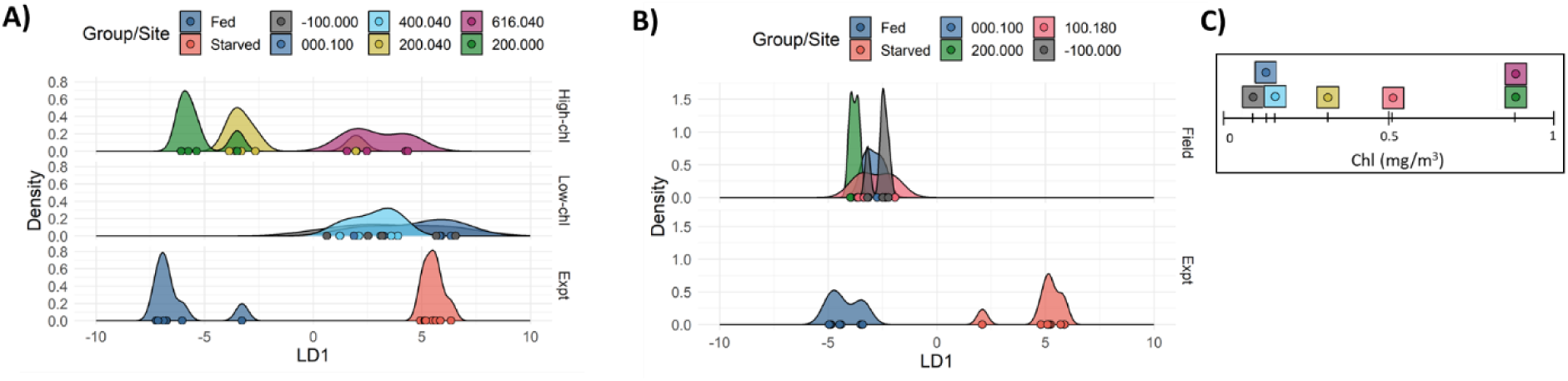
Comparison of transcriptomic patterns from field stations and starvation experiments using discriminant analysis of principal components for (C) *C. acutus* and (D) *C. propinquus*. LD1 is the linear discriminant axis that maximizes variation between Fed and Starved samples (“Expt”, bottom plot). (C) Stations 200.xxx overlap with Fed samples, and low-chlorophyll *a* stations overlap with Starved samples. The three highest-chl *a* (top) and three lowest-chl *a* stations (middle) are shown separately for clarity. (D) All *C. propinquus* field stations cluster with Fed samples. (E) Field stations ordered according chl *a* concentration (averaged over top 200 m).

For *C. acutus,* we noted that gene expression changes between station groups broadly reflected the experimental starvation response. The correlation between log2(fold change) of DE genes in the starvation experiment and log2(fold change) between station groups was ρ=0.79, slightly stronger than the correlation with chl *a* itself. Of 3687 DEGs in the starvation experiment, 2527 (69%) were also DE between station groups. For >99% of these (2520/2527), stations 200.xxx had gene expression signatures consistent with fed animals (Table 1). This overall similarity was also reflected in the WGCNA analysis, as the Red, Salmon, and Turquoise modules were upregulated in fed animals and at stations 200.xxx, and the Black, Magenta, and Yellow modules were downregulated in fed animals and at stations 200.xxx (Fig. 6).

For *C. propinquus,* we identified fewer DEGs within both the field study and experiment. Of 1186 DEGs from the starvation experiment, only 27 were also DE in any comparison among stations. Of these, most (20) were upregulated at 200.000 and in the fed group. WGCNA showed some concordance between field chl *a* and experimental starvation, as the Ivory, Darkorange and Purple modules were downregulated in fed animals and at station 200.000. In contrast, the Royalblue module was downregulated in starved animals and station 000.100 but upregulated at

200.000 (Fig 7). Thus, although *C. propinquus* had a weaker association between field chl and starvation overall, animals at site 200.000 appear to be somewhat better fed than other sites, because they have reduced expression of starvation-associated WGCNA modules and higher expression of some lipid-associated genes.

### Putative biomarkers of feeding status

We next sought to identify genes that might reliably indicate feeding status by selecting genes with the most significant changes in expression in both the starvation experiment and the field study. In the field study, we selected genes that were DE at stations with signatures of favorable feeding conditions: 200.xxx for *C. acutus* and 200.000 for *C. propinquus*. After applying filtering criteria based on expression, effect size, and significance in both studies (see Methods), we identified 183 such genes in *C. acutus* and 25 in *C. propinquus*. Two homologous genes were found in both lists: a vitellogenin, which had one copy in *C. propinquus* and two paralogs in *C. acutus*, and a methyltransferase, also with one copy in *C. propinquus* and two copies in *C. acutus*.

In *C. acutus*, 182 of 183 putative biomarkers were downregulated in starved animals; the upregulated biomarker was annotated as an enoyl-CoA hydratase, which is involved in fatty acid catabolism. Downregulated biomarkers included several genes with roles in amino acid catabolism (an L-threonine 3-dehydrogenase, a proline dehydrogenase, a phenylanine/tryptophan hydroxylase), a gene that catalyzes a step in fatty acid degradation (inorganic pyrophosphatase; Chandel, 2021), and gene with key roles in lipid synthesis (desaturases and an elongase; these are among the “lipid-associated genes” in Table 2). In *C. propinquus*, 6 biomarkers were upregulated, including a CREB-like gene, triacylglycerol lipase, and phosphatidate phosphatase. This last gene catalyzes a rate-limiting step in triglyceride synthesis and also has roles in lipid signaling (Brindley, 1984; Chandel, 2021). Downregulated biomarkers in *C. propinquus* included several possible hemerythrins (oxygen-binding proteins) and a tricarboxylate transfer protein.

## Discussion

Field-based studies of gene expression have the potential to identify environmental drivers of physiological variation among marine organisms. However, field studies can be challenging to interpret because environmental factors are confounded with many other (possibly unobserved) variables. One promising approach is to pair field studies with experiments to determine how specific factors, such as food availability, contribute to physiological variation *in situ*. In this study, we assayed patterns of gene expression and other physiological metrics across stations within the PAL-LTER sampling grid for two key species of Southern Ocean zooplankton, *Calanoides acutus* and *Calanus propinquus*. By comparing field data with starvation experiments conducted on the same research cruise (Berger et al., 2023), we provide evidence that field chl *a* concentration generally reflects local food availability on a timescale of ∼1 week, and that variation in food availability is a major factor associated with copepod physiological variation among sites, especially for the largely herbivorous *C. acutus*. We also identify candidate biomarker genes whose expression reliably indicates feeding status. While we have identified some similarities in the physiological patterns between the two species, we found female *C. acutus* to be more sensitive than *C. propinquus* to short-term variation in food availability and to exhibit more spatial variation in physiological condition. These properties are consistent with a strong reliance of *C. acutus* on continued feeding on phytoplankton to support egg production.

### Strong separation based on feeding condition between sites for an herbivorous grazer

For *C. acutus*, we focused our analysis on two stations that formed a distinct cluster in the gene expression analysis. Stations 200.00 and 200.040 both had moderate-to-high chl *a* in the upper 200m, and copepods at these sites were actively feeding and had elevated citrate synthase activity, indicating elevated aerobic metabolism. The stations are both located within Marguerite Bay, an area of high productivity that sustains large zooplankton populations and isan important foraging ground for penguins, whales and other large predators (Casanovas et al., 2015; Deibel & Daly, 2007; Rozema et al., 2017; Siegel et al., 2013). No *C. acutus* from any other site was observed with food in its gut, despite one other site having comparably high *chl a*. There was strong concordance between starvation-response genes determined experimentally (Berger et al., 2023) and genes that were differentially expressed between station groups, indicating that separation in gene expression space was largely driven by differences in feeding status. High expression of lipid synthesis enzymes at these stations suggests active lipid synthesis in food-replete conditions, while upregulation of vitellogenins and other genes related to reproduction suggests partitioning of lipids into reproductive capacity.

These results are consistent with studies of *Neocalanus flemingeri* in the Gulf of Alaska, which noted differences in metabolic gene expression associated with chlorophyll variation across sites (Roncalli et al., 2019) and between years (Roncalli et al., 2022). This suggests that chlorophyll levels are a primary factor contributing to gene expression differences among field sites for herbivorous copepods. Like *N. flemingeri*, *C. acutus* relies on short phytoplankton blooms to mature, reproduce, and perform diapause.

Although there was a strong overall association between field chl *a* and experimental starvation response-genes for *C. acutus*, one high-chl *a* site (616.040) grouped with low-chl *a* sites in gene expression analyses (Figs. 1 & 9). This site had a uniformly warm water column in the upper 200m and an intense, but shallow, surface chl *a* maximum; chl *a* maxima at other sites were deeper. No animals sampled from this site had food in their gut, samples were more similar to starved than fed experimental samples, and animals had low citrate synthase activity, indicating poor feeding conditions for *C. acutus* despite the high chl *a* concentration. One possible explanation is that this site had a distinct phytoplankton community: large calanoid copepods feed preferentially on diatoms compared to, for instance, the haptophyte *Phaeocystis spp.,* which is a lower-quality food source (Head & Harris, 1994; Turner et al., 2002). Although overall chl *a* levels may be a useful proxy for food availability for grazers, the specific local abiotic and biotic factors, including phytoplankton community composition, determine feeding favorability.

### Lesser variation for an omnivore

*C. propinquus* physiology and gene expression exhibited less variation among sites than *C. acutus*. While this may be partially attributed to the smaller sample size for *C. propinquus*, the more omnivorous diet of *C. propinquus* may also reduce its sensitivity to short-term variation in phytoplankton abundance. Consistent with this, *C. propinquus* had a much weaker correlation between experimental starvation and field *chl a* than *C. acutus*. Nonetheless, some starvation-response genes were associated with field chl *a*, and one high-chl *a* site (200.000) had elevated expression of lipid synthesis genes (elongases and desaturases) and vitellogenins, suggesting favorable feeding conditions. This was also a “favorable” site for *C. acutus*, suggesting that the same favorable feeding conditions apply to both species. Therefore, chl *a* levels seem to drive some physiological variation in *C. propinquus*, but to a lesser extent than *C. acutus*.

None of the *C. propinquus* field samples grouped with starved experimental samples in our DAPC analysis, suggesting that none of the field samples had reached a comparable state of food deprivation. Indeed, animals were visibly feeding on algae at all sites. In contrast, DAPC of *C. acutus* suggests that the field samples spanned the entire range of nutritional conditions captured by the starvation experiment. Although *C. propinquus* might genuinely exhibit less physiological variation among sites, sampling over a larger range of stations may uncover additional variation in this species.

### Chlorophyll *a* is a predictive factor in copepod physiology as well as population dynamics

Long-term monitoring within the WAP region has shown that interannual variation in chl *a* concentration is strongly associated with variation in *C. acutus* abundance (Gleiber, 2014), as well as abundance of total copepods. Concordant with this, we observed a spatial correlation between chl *a* and *C. acutus* physiological condition. Gleiber (2014) also found an association between *C. propinquus* abundance and chl *a* lagged by 1 year. This lagged relationship may reflect the reduced dependence of *C. propinquus* on phytoplankton and greater capacity for lipid storage; these traits may underlie our observations of a weaker relationship between chl *a* and *C. propinquus* physiology. While chl *a* is associated with interannual variation in the abundances of both species, and contributes to spatial variation in physiological condition, we did not find a spatial relationship between chl *a* and the abundance of either species. Despite the apparent favorability of site 200.000 for both species (and site 200.040 for *C. acutus*), *C. acutus* and *C. propinquus* were much more abundant farther south. This disconnect may be driven by small-scale mismatches between the phytoplankton and copepod populations, differential losses from predation, or reduced recruitment and retention due to currents.

From an evolutionary perspective, advection limits capacity of pelagic zooplankton for local adaptation, and many species have little population structure even on basin scales (Choquet et al., 2019). In such species, physiological plasticity may be more important than local adaptation for determining species distributions and population dynamics in the warming ocean. In the WAP, warming conditions and increased stratification are expected to lead to decreased production and biomass of large phytoplankton, especially in the northern region (Ferreira et al., 2020; Gleiber et al., 2016; Schofield et al., 2018; Venables et al., 2013). For a species such as *C. acutus* that depends on seasonal phytoplankton blooms, these changes will certainly result in shifts in spatial patterns of habitat suitability, as well as possible phenological mismatches. *C. propinquus* may be more robust to these changes due to its greater degree of omnivory, but nonetheless needs phytoplankton to feed and reproduce optimally. The favorability of local feeding conditions affects the fitness and reproductive capacity of a population. Ultimately, spatial variation in fitness, together with advection, may contribute to regional differences in recruitment and source-sink dynamics and shape long-term population distributions.

### Feeding status biomarkers differ between species

Although many homologous genes respond similarly to starvation in both species (Berger et al., 2023), we identified almost no homologous “biomarker” genes consistently associated with feeding status in both the field study and starvation experiment. Identification of such genes may have been hampered by the limited range of conditions captured in our sampling of *C. propinquus*. Nonetheless, vitellogenins were strongly downregulated in both species with starvation and at low-chl *a* stations. Since vitellogenins are often highly expressed and sensitive to food availability, they may be particularly reliable indicators of recent feeding status. Vitellogenin expression also has a clear biological interpretation—females increase their reproductive investment when they are better fed. Unfortunately, this biomarker is largely restricted to reproductively mature (or maturing) females. Considering each species separately, genes involved in lipid metabolism appear to be reasonable indicators of metabolic condition, including desaturases and elongases (*C. acutus*) and lipases (*C. propinquus*).

Overall, our results lend confidence to interpretations of field-based RNA-seq studies of pelagic copepods. Much of the spatial variability in *C. acutus* gene expression could be attributed to differences in feeding status, and even the omnivorous *C. propinquus* exhibited some variation attributable to food availability. We identified highly-expressed genes associated with both field chl *a* and starvation responses, which may be used to infer relative differences in recent feeding status of these species in future field studies. Researchers should consider not only gross measures of abundance but also spatial variation in gene expression and other physiological metrics. These integrated studies will ultimately improve our capacity to understand and predict how copepod populations respond to changing conditions in the WAP ecosystem and other regions.

## Author Contributions

Designed research: AMT, DKS, CAB; Performed field sampling: AMT, DKS; Performed laboratory analyses: AMT; Analyzed data: CAB, AMT; Wrote the paper CAB, AMT. All authors reviewed and edited the final manuscript.

## Acknowledgements

We thank the officers, crew and technical staff of the ARSV *Lawrence M. Gould*; Joe Cope, Dr. Patricia Thibodeau, and other members of the Steinberg Lab for copepod sampling and other assistance at sea; and Nancy Copley and Adrienne Jones for assistance in photo analysis. Funding for this project was provided by the National Science Foundation Office of Polar Programs (Grants OPP-1746087 to AMT, and OPP-1440435 and OPP-2026045 to DKS).

## Data Accessibility

Raw RNA-seq data have been uploaded to the NCBI Sequence Read Archive (SRA), Bioprojects PRJNA757455 (*Calanoides acutus*) and PRJNA669816 (*Calanus propinquus*). R code used for analysis is available at https://github.com/caberger1/Copepod_starvation_scripts.

