## Supplemental File 1 for "Nutritional condition drives spatial variation in physiology of Antarctic lipid-storing copepods"

MOLECULAR ECOLOGY

### Table of Contents:

| | |
|---|---|
| **Supplemental Tables** | Page 2 |
| **Supplemental Figures** | Page 6 |

MOLECULAR ECOLOGY

Table S1: Locations of stations sampled for copepod physiological measurements (transcriptomics = T; citrate synthase = CS, X = neither) and/or species abundance (individuals per 1000 m$^3$). Species names: *Calanoides acutus, Calanus propinquus, Rhincalanus gigas*

| Station | Latitude S | Longitude W | Date | Physiology | | Abundance | | |
|---|---|---|---|---|---|---|---|---|
| | | | | *C. acutus* | *C. propinquus* | *C. acutus* | *C. propinquus* | *R. gigas* |
| -100.100 | 68°49.171’ | 77°05.016’ | 01/23/19 | T, CS | T, CS | 4516 | 30 | 133 |
| -100.160 | 68°22,875’ | 77°55.286’ | 01/25/19 | X | X | 1568 | 6.7 | 628 |
| 000.100 | 68°16.583’ | 75°07.077’ | 01/22/19 | T, CS | T. CS | 1201 | 23 | 45 |
| 100.180 | 67°09.441’ | 74°28.573’ | 01/21/19 | X | T | 962 | 28 | 203 |
| 200.-040 | 68°2.271’ | 69°18.376 | 01/17/19 | X | X | 532 | 1.1 | 24 |
| 200.000 | 67°46.319’ | 69°56.667’ | 01/17/19 | T, CS | T | 295 | 9.0 | 164 |
| 200.040 | 67°30.664’ | 70°35.377’ | 01/14/19 | T, CS | X | 1051 | 12 | 83 |
| 400.040 | 66°15.237’ | 67°20.194’ | 01/10/19 | T | X | 358 | 9.5 | 42 |
| 500.100 | 65°14.029’ | 66°46.558’ | 01/31/19 | X | X | 235 | 7.7 | 51 |
| 600.100 | 64°34.516’ | 65°20.483’ | 01/05/19 | X | X | 81 | 4.2 | 33 |
| 616.040 | 64°50.101’ | 64°09.763 | 02/04/19 | T | X | 15 | 3.2 | 46 |

Table S2. *Calanoides acutus* observations of food in gut and reproductive condition. Note that sample sizes reflect the photographs of sufficient quality to evaluate, which differed depending on the observation type.

| **Station** | **Food in Gut** | **Reproduction**[1] |
|---|---|---|
| -100.100 | Yes: 0, No: 10, N: 10 | Min: 12, Mod: 8, WD: 2, N: 22 |
| 000.100 | Yes: 0, No: 12, N: 12 | Min: 18 Mod: 3, WD: 4, N: 25 |
| 200.000 | Yes: 7, No: 5, N: 12 | Min: 16, Mod: 13, WD: 6, N: 35 |
| 200.040 | Yes: 8, No: 23, N: 31 | Min: 14, Mod: 5, WD: 10, N: 29 |
| 400.040 | Yes: 0, No: 4, N: 4 | Min: 1, Mod: 0, WD: 1, N: 2 |
| 616.040 | Yes: 0, No: 9, N: 9 | Min: 4, Mod: 4, WD: 4, N: 10 |
| **Station Group** | **Food in Gut** | **Reproduction**[1] |
| 200.000 & 200.040 | Yes:15, No: 28, N: 43 | Min: 30, Mod: 18, WD: 16, N: 64 |
| Others | Yes: 0, No: 35, N: 35 | Min: 35, Mod: 15, WD: 9, N: 59 |

[1] Reproductive categories: Min = Minimal egg development, Moderate egg development = Moderate , WD = Well-developed eggs (“N” denotes sample size)

MOLECULAR ECOLOGY

**Table S3:** ANOVA and post-hoc statistics for *Calanoides acutus* citrate synthase measurements

Table SB1: Welch's ANOVA results for physiological data

| **Measurement** | **d.f.[1]** | **F** | **p-value** |
|---|---|---|---|
| Citrate synthase | 3, 12 | 8.07 | <0.001 |

| **Measurement** | **Contrast station pair[1]** | **T** | **d.f.** | **p-value** |
|---|---|---|---|---|
| Citrate synthase | 200.000 vs 000.100 | 5.242 | 6 | 0.001 |
| | 200.000 vs -100.100 | 5.750 | 6 | 0.001 |
| | 200.040 vs 000.100 | 3.920 | 6 | 0.006 |
| | 200.040 vs -100.100 | 4.427 | 6 | 0.003 |

MOLECULAR ECOLOGY

Table S4: *Calanoides acutus* Gene Ontology Terms enriched in the gene expression contrast between stations 200.040 and 200.000 (200.xxx) and all other stations. Selected metabolic categories shown; for full results see Supp. File 2. Italicized terms were collapsed during REVIGO summarization.

| Category | Up in 200.xxx | Down in 200.xxx |
|---|---|---|
| Lipid Metabolism | regulation of lipid metabolic process, regulation of lipid biosynthetic process | None |
| Protein Metabolism | histone modification, histone deacetylation, regulation of histone methylation, positive regulation of cysteine-type endopeptidase activity involved in apoptotic process | cytoplasmic translation, regulation of translation, proteolysis, protein ubiquitination, mitochondrial translation |
| Response to Stress | negative regulation of oxidative stress-induced cell death | DNA repair, *cellular response to stress* |
| Reproduction | meiotic gene conversion, mating behavior, *ovarian follicle cell development, positive regulation of reproductive process* | None |

Table S5. *Calanus propinquus* observations of food in gut and reproductive condition. Note that sample sizes reflect the photographs of sufficient quality to evaluate, which differed depending on the observation type.

| Station | Food in Gut | Reproduction[1] |
|---|---|---|
| -100.100 | Yes: 14, No: 3, N: 17 | Min: 8, Mod: 3, WD: 3, N: 14 |
| 000.100 | Yes: 10, No: 2, N: 12 | Min: 6 Mod: 3, WD: 3, N: 12 |
| 100.180 | Yes: 6, No: 12, N: 18 | Min: 5, Mod: 3 WD: 10, N: 18 |
| 200.000 | Yes: 5, No: 3, N: 8 | Min: 1, Mod: 1, WD: 3, N: 5 |

[1] Reproductive categories: Min = Minimal egg development, Moderate egg development = Moderate , WD = Well-developed eggs ("N" denotes sample size)

MOLECULAR ECOLOGY

Table S6: *Calanus propinquus* Gene Ontology Terms enriched in the gene expression contrast between each station and the mean of all other stations. Selected metabolic categories shown: lipid- and reproduction-associated genes, and enriched GO terms associated with lipid metabolism (L), response to stress (S) and reproduction (R). For full results see Supp. File 3.

| Station | Upregulated | Downregulated |
|---|---|---|
| 200.000 | L: very long-chain fatty acid metabolic process, regulation of steroid metabolic process<br>P: regulation of histone methylation, histone H3-K9 methylation<br>S: response to ischemia, S/R: double-strand break repair involved in meiotic recombination<br>R: positive regulation of meiotic cell cycle, | P: negative regulation of histone H3-K3 methylation, translation<br>S: positive regulation of double-strand break repair via nonhomologous end joining |
| 100.180 | R: border follicle cell migration | |
| 000.100 | L/P: dolichol-linked oligosaccharide biosynthetic process;<br>P: translation, regulation of translation, proteasomal ubiquitin-independent protein catabolism, protein modification by small protein conjugation or removal<br>S: regulation of transcription from RNA polymerase II promoter | |
| -100.100 | P: regulation of protein phosphorylation<br>S: response to laminar fluid shear stress | L: regulation of steroid metabolic process<br>P: translational initiation, histone deacetylation, regulation of histone methylation<br>S: response to hypoxia, response to ischemia, cellular response to DNA damage stimulus, regulation of oxidative stress-induced cell death;<br>R: multi-organism reproductive process |

MOLECULAR ECOLOGY

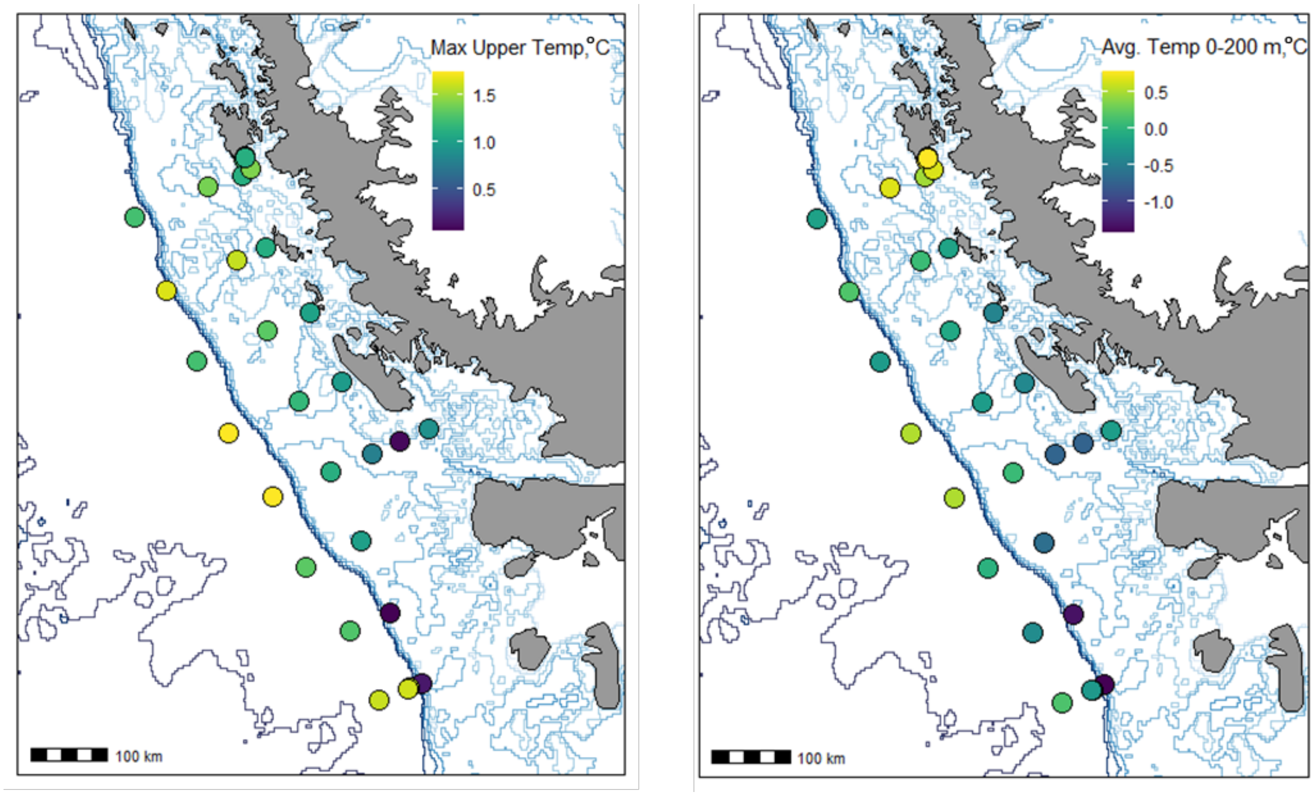

Figure S1. Maximum (left) and average (right) temperature (°C) within the upper 200 m depth.

MOLECULAR ECOLOGY

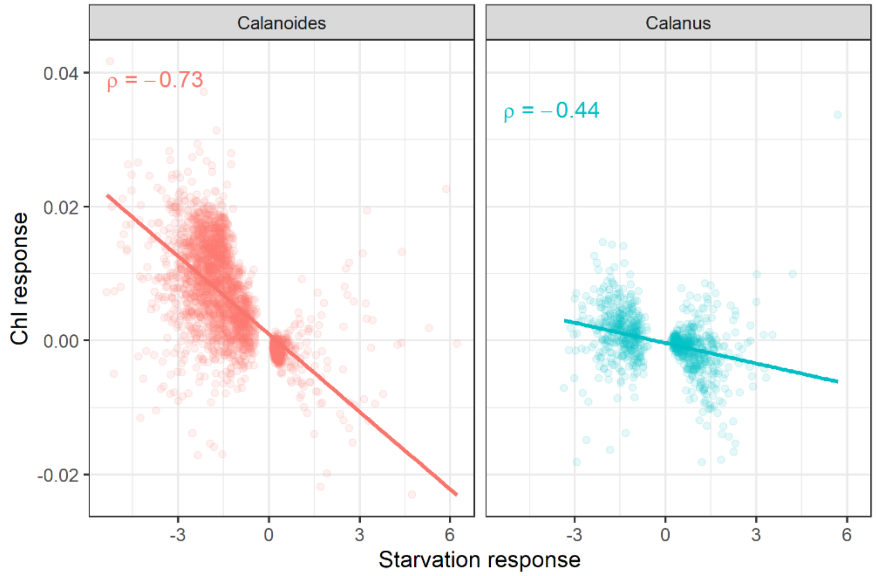

Figure S2: Correlation (Spearman's) between the fold-change associated with chlorophyll a concentration in the field (y-axis) and the fold change associated with starvation in *Calanoides acutus* (left) and *Calanus propinquus* (right).
